# *In situ* transcriptome characteristics are lost following culture adaptation of adult cardiac stem cells

**DOI:** 10.1101/359497

**Authors:** Taeyong Kim, Oscar H. Echeagaray, Bingyan J Wang, Alexandria Casillas, Kathleen M. Broughton, Bong-Hyun Kim, Mark A Sussman

## Abstract

Regenerative therapeutic approaches for myocardial diseases often involve adoptive transfer of stem cells expanded *ex vivo*. Prior studies indicate that cell culture conditions affect functional and phenotypic characteristics, but relationship(s) of cultured cells derived from freshly isolated populations and the heterogeneity of the cultured population remain poorly defined. Functional and phenotypic characteristics of adoptively donated cells will determine outcomes of interventional treatment for disease, necessitating characterization of the impact that *ex vivo* expansion has upon isolated stem cell populations. Single-cell RNA-Seq profiling (scRNA-Seq) was performed to determine consequences of culture expansion upon adult cardiac progenitor cells (CPCs) as well as relationships with other cell populations. Bioinformatic analyses reveal loss of identity marker genes in cultured CPCs while simultaneously acquiring thousands of additional genes. Cultured CPCs exhibited decreased transcriptome variability within their population relative to their freshly isolated cells. Findings were validated by comparative analyses using scRNA-Seq datasets of various cell types generated by multiple scRNA-Seq technology. Increased transcriptome diversity and decreased population heterogeneity in the cultured cell population relative to freshly isolated cells may help account for reported outcomes associated with experimental and clinical use of CPCs for treatment of myocardial injury.

## Introduction

Stem cell therapy is a promising approach for mitigating pathological diseases such as heart failure, with cell populations derived from diverse origins proposed for autologous as well as allogeneic cell therapy^1-3^. The presumption that donor cells retain essential characteristics derived from their original identity during *in vitro* expansion important to enhance regeneration has led to isolation of cardiac progenitor cells (CPCs) subjected to culture for expansion prior to reintroduction. Multiple donor cell types have been tested for basic biological characteristics and efficacy, with widely varying isolation and adoptive transfer methods^4,5^. For example, CPCs used in clinical trials for cardiac repair are isolated and cultured using varying and unstandardized protocols^6-9^. Transcriptome profiling of cultured CPCs using varying isolation methods showed surprisingly high similarity^10^, possibly accounting for consistently modest functional improvement outcomes in the myocardium regardless of cell type^3^. However, bulk RNA sample profiling of cultured CPCs in prior studies masks population heterogeneity inherent to freshly isolated CPCs^11^. Therefore, understanding the consequences and impact of culture expansion upon the transcriptome at the single cell level is essential to optimize and advance approaches intended to improve efficacy of stem cell-based cardiac regenerative therapy.

Transcriptome profiling of freshly isolated CPCs is challenging due to low yields of resident adult stem cells, with very limited transcriptome information on primary isolates of other stem cells^12-15^. Implementation of single-cell RNA-Seq (scRNA-Seq) allows for transcriptional profiling of low cell numbers as well as revealing population heterogeneity. Technical aspects of scRNA-Seq tend toward choosing between transcriptome depth with limited number of cells versus massively parallel sequencing using hundreds to thousands of cells with shallower transcriptome coverage. Recent advances in massively parallel scRNA-Seq demonstrate the capability to maximize number of single cells captured per sample while still capturing primary characteristics of transcriptome variation^11,16,17^. Unfortunately, the relatively recent advent of massively parallel scRNA-Seq has yet to produce the range and depth of scRNA-Seq datasets acquired using Smart-Seq2 technology that is limited by small population samples^18^. Therefore, a combination of both scRNA-Seq approaches involving Smart-Seq2 as well as massively parallel transcriptome profiling was used to determine the transcriptome identity and population heterogeneity of CPCs either as freshly isolates versus their cognate cultured counterparts. Based on the scRNA-Seq data analysis comparing freshly isolated cells and cultured cells, we identified common and global transcriptome alterations consequential to *in vitro* expansion. Findings reveal that isolation and *in vitro* expansion of CPCs selects for transcriptional profiles of uniform composition resulting in loss of *in situ* characteristics as well as population heterogeneity. The consequences of this transcriptional drift and homogenization of cellular phenotypes offers fundamental biological insight regarding the basis for consistently modest efficacy of CPC-based cell therapy and prompts reassessment of the rationale for tissue-specific stem cell sources.

## Results

### Transcriptome drift of freshly isolated CPCs following short term culture

Transcriptional profiling was performed using freshly isolated cells and their *in vitro* derivatives to reveal consequences of short term culture. Population characteristics were revealed by scRNA-Seq using the 10x Chromium platform. Freshly c-kit+ / Lin-CPCs were prepared for profiling in tandem with cultured CPC populations expanded under standard conditions^19^ for five passages (Fig1a). Both fresh and cultured CPC scRNA-Seq datasets were mapped to mouse genome, aggregated using Cell Ranger v2.0 (10X Genomics), and unsupervised clustering performed using Seurat R package^20^ (Fig. 1b). Separation between fresh versus cultured CPCs clusters was clearly demonstrated by t-SNE plot^21^, revealing divergence of transcriptome between these two cell populations based upon spatial distance. Robustness of ‘clear separation’ between fresh cells and cultured cells was tested with multiple different parameter settings as previously reported for fresh murine heart cell isolates.^11^ Clustering is remarkably robust regardless of parameter setting for t-SNE plotting (Fig. 1c) such as perplexity or the number of principal components (PC). Clustering results reflect differences between fresh and cultured cells (Fig. S1) and principal component analysis (PCA) also showed distinctive clustering in consistent with t-SNE results (Fig. S2). Collectively, these findings reveal substantial transcriptome divergence between fresh versus cultured CPCs at the single cell level.

**Figure 1.**
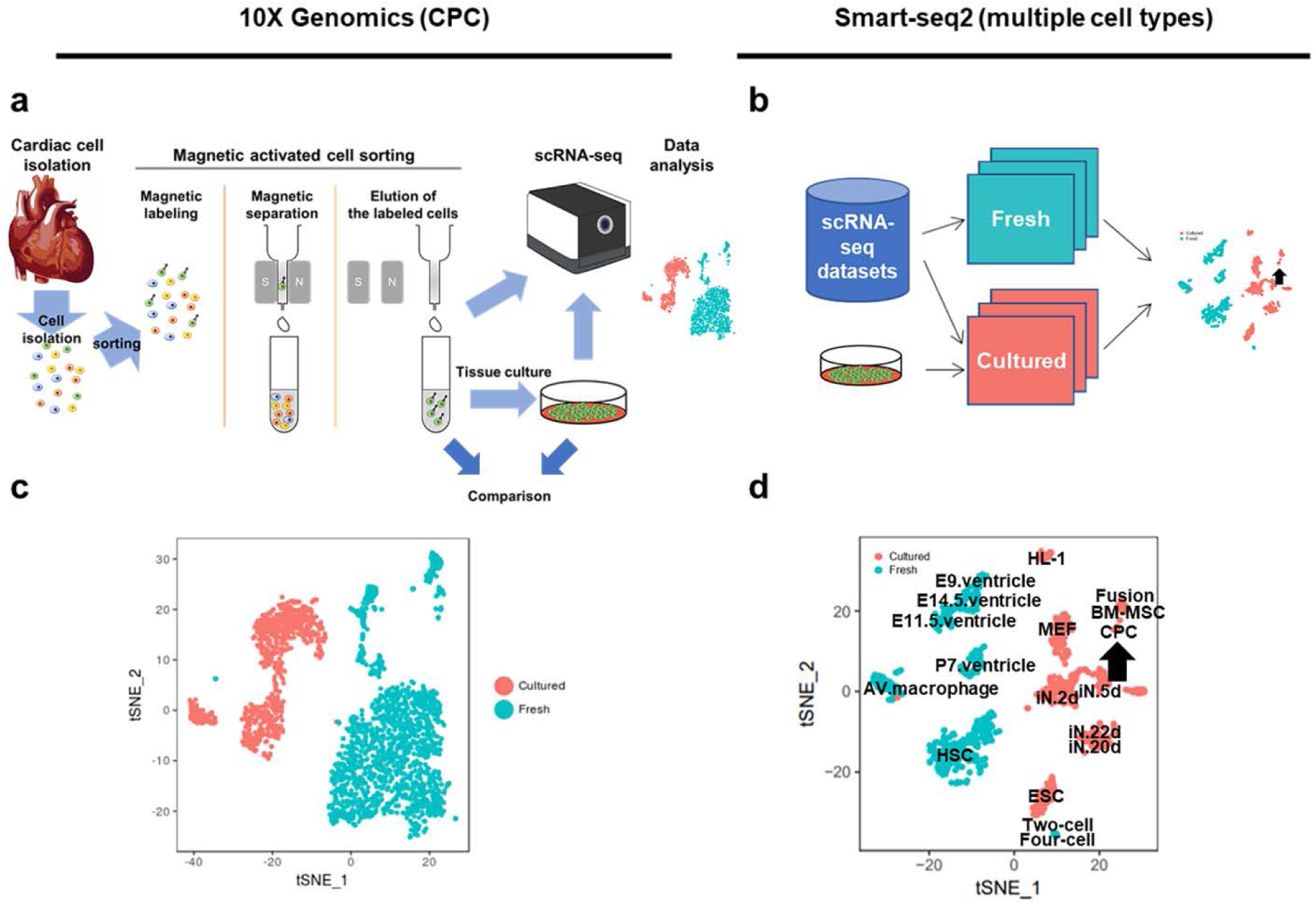
Transcriptomic differences between freshly isolated and cultured cells revealed by scRNA-Seq. (a-b) Workflow of scRNA-Seq of CPCs using 10X Genomics (a) or dataset of multiple cell types using Smart-Seq2 technology (b). (c-d) t-SNE plots show freshly isolated cells cluster separately from cultured cells using 10X Genomics dataset (c) or Smart-seq2 dataset (d). Arrow indicates CPCs prepared and sequenced in this study relative to single-cell datasets downloaded from public databases (see supplemental table for details). CPC, cardiac progenitor cell; BM.MSC, bone marrow-derived mesenchymal stromal cell; MEF, mouse embryonic fibroblast; ESC, embryonic stem cell; iN, induced neuronal cell; HL-1, atrial cardiomyocyte cell line; Fusion, Fusion cell of BM-MSC and HL-1; AV macrophage, atrioventricular node macrophage; E9, E11.5, E14.5, embryo 9, 11.5, 14.5 days, respectively; P7, postnatal 7 days; HSC, hematopoietic stem cell

Analyses were broadened to determine consequences of culture upon additional cell populations upon CPCs relative to other stem cell types. Deeper transcriptome coverage for CPCs was achieved using Smart-Seq2 technology as further validation of 10X Genomics results (Fig. 1) that provides wider population-based coverage but possesses limited resolving power for distinguishing marginally different cell types limited due to shallower sequencing depth (50,000~100,000 reads per cell). Meta-analysis of multiple scRNA-Seq datasets including both variety of fresh cells and various cultured cells was performed ^22-28^ (Supplementary table) using single-cell datasets (around 1,500 cells) generated by Smart-Seq2.^18^ Processing of datasets was performed comparably to initial CPC analyses with mapping to mouse genome and clustering analysis using Seurat R Package. Unsupervised clustering showed similar clear separation between fresh and cultured cells in Smart-Seq2 datasets (Fig. 1d) comparable to findings with 10X Genomics (Fig. 1a). Normalization of downloaded Smart-Seq2 scRNA-Seq datasets for sequencing depth prior to comparison to exclude effects of sequencing depth variation was also performed, downsampling to 500,000 reads per cell by random selection of raw reads from fastq files and re-analyzed. Comparable trends of separation between fresh versus cultured cells remained in meta-analysis of downsampled data (Fig. S3), reinforcing the conclusion that transcriptomic alteration by tissue culture is a shared phenomenon regardless of cell type and consistent between the two scRNA-Seq methodologies of 10X Genomics and Smart-Seq2.

### Subpopulations revealed by clustering analyses of freshly isolated and cultured CPCs

Three distinct clusters exist in freshly isolated CPCs on the t-SNE plot revealed after dimensionality reduction followed by unsupervised clustering (Fig. 2a-b), two of which identify as endothelial cells and hematopoietic cells based on the gene set analysis (Fig. S4a). A third cluster represents a heterogeneous cell population including smooth muscle cells, pericytes, and two different fibroblast subpopulations (FigS4b-c) based upon marker expression of cardiac cells.^11^ This unbiased third cluster consisting of fibroblasts, smooth muscle cells, and pericytes was collectively categorized as ‘cardiac interstitial cells’ (CICs). Cultured cells were divided into two groups, ‘CultureA’ and ‘CultureB’ through unsupervised clustering (Fig. 2a and b). Gene set analysis using a set of differentially expressed genes (DEGs) between CultureA and CultureB was performed to identify cell type of the two cultured cell groups. Genes enriched in CultureA primarily include cell adhesion-related genes, while genes of CultureB cluster include respiratory energy metabolism-related genes (Fig. S5a-d). Although ClusterA and Cluster B are different through gene set analysis, transcriptomic signature was lacking since no cell type-specific gene set was detected. Additionally, fresh CPC marker expression^29^ levels were comparatively low in cultured CPCs (Fig. S5f). Each cluster of freshly isolated cell possesses a unique set of DEGs not detected in other clusters, while the two clusters of cultured cell share their DEGs (Fig. 2c). In summary, sub-populations were readily identifiable in freshly isolated CPCs, but not evident in cultured CPCs.

**Figure 2.**
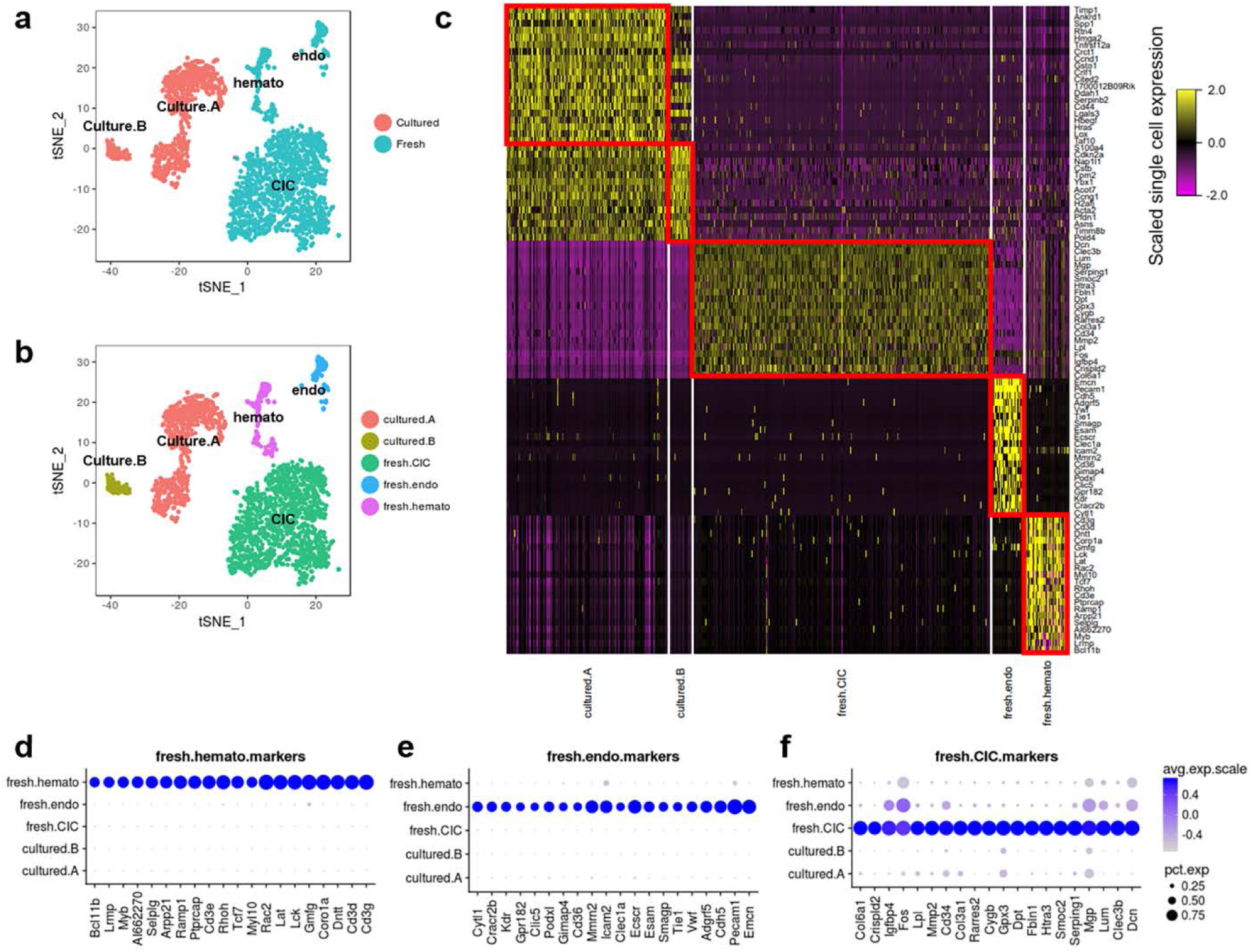
Loss of identity markers following *ex vivo* expansion. (a and b) t-SNE plots show single cells labeled by environments (a) or cell types (b). Cell clusters are identified by expression level of marker genes and gene set analysis as shown in Fig. S4 and Fig. S5. (c) Heatmap representing top 20 marker genes of each cluster. (d-f) Dotplots representing top 20 marker genes of each cluster. Marker genes were identified in an unbiased fashion blind to known cell type markers. Individual dots are sized to reflect the proportion of cells of each type expressing the marker gene and colored to reflect the mean expression of each marker gene across all cells as indicated in the key.

### Loss of identity markers and enhanced cell proliferation consequential to culture expansion

The relationship between fresh versus cultured CPCs was assessed for expression of marker genes between clusters. Identifier marker genes for fresh cell clusters were substantially down-regulated in cultured cells (Fig. 2d-f; Fig. S5e). Marker transcript identity for cultured cells revealed protein metabolism and cell cycle pathways were enriched in the 10X Genomics cultured dataset assessed by searching for upregulated genes using gene set analysis with gene ontology (GO) terms. Elevated expression of transcripts associated with protein synthesis and cell proliferation is consistent with biological selective pressures of *in vitro* expansion (Fig. 3a). Proliferation-related pathways (such as ‘mesenchymal cell proliferation’) were detected as a common pathway between 10X Genomics and Smart-Seq2 data (Fig. 3b). Expression patterns of all cell cycle (CC) -related genes was assessed in both datasets^30^ to confirm the result of gene set analysis at individual gene level. Multiple G1/S phase- and G2/M phase-specific genes were commonly upregulated in cultured cells (Fig. 3c-f) including aurora kinases (Aurka and Aurkb), cyclin B2 (Ccnb2), centromere proteins (Cenpa, Cenpe and Cenpf) and cyclin-dependent kinase 1(Cdk1) in both 10X Genomics and Smart-Seq2 datasets. Based upon expression level of CC genes, cell cycle scores were calculated^30^ and cell cycle stages were estimated (Fig. 3g-j). A higher G2/M ratio was present in cultured cells indicative of proliferative state (average G2/M ratio: 4.8% and 29.0% in fresh and cultured CPCs, respectively; 9.3% and 43.7% in fresh and cultured cells from meta-analysis, respectively). Collectively, cultured cells are characterized by loss of original identity markers and a transcriptome profile characterized by up-regulated cell proliferation genes.

**Figure 3.**
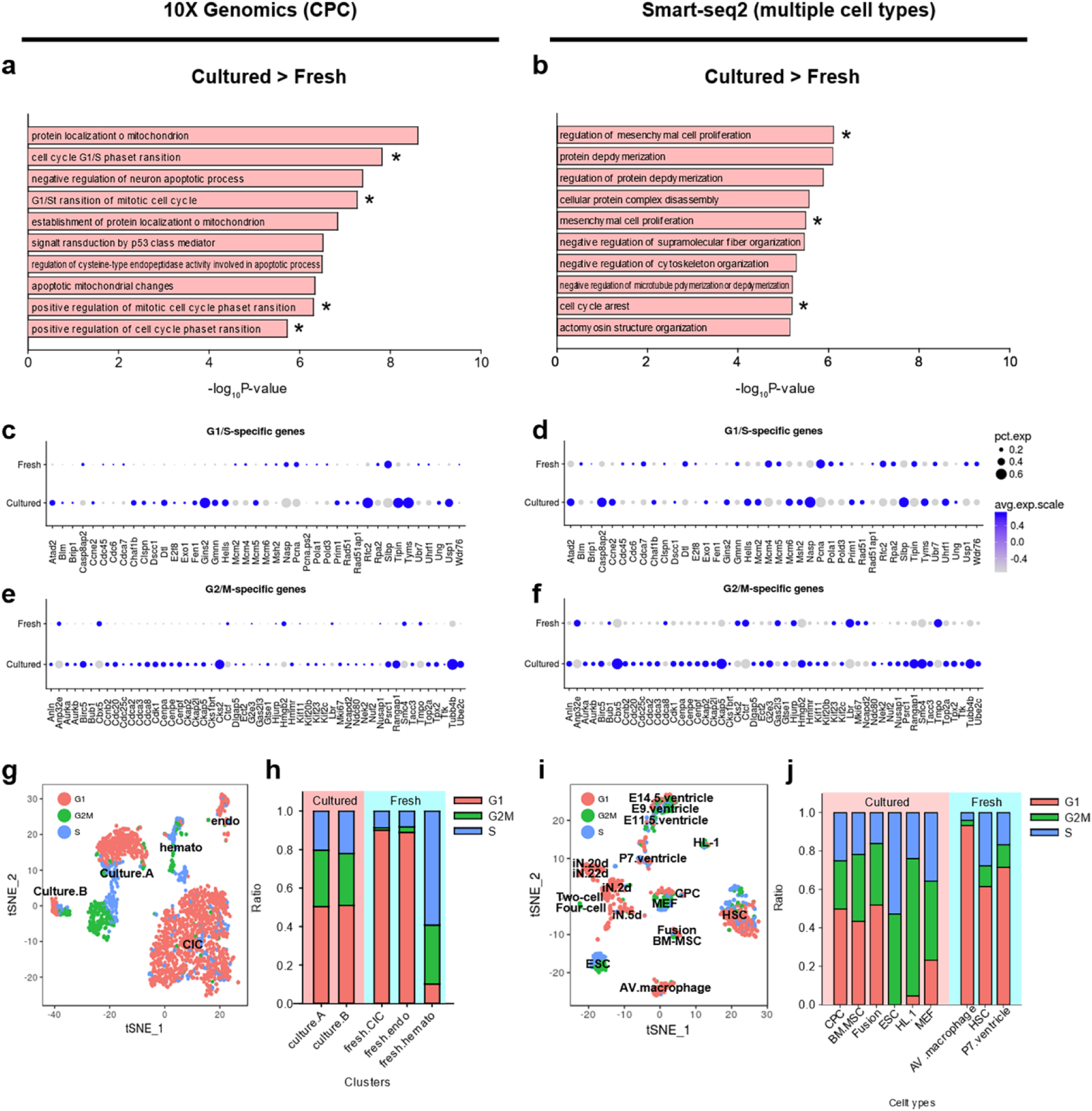
Culture expansion increases expression of cell proliferation-related genes. (a and b) Bar graph represents top 10 GO terms up-regulated in cultured cells revealed by gene set enrichment analysis from 10X Genomics (a) or Smart-Seq2 data analysis (b). Asterisks represent cell proliferation-related GO terms commonly observed in both 10X Genomics and Smart-Seq2. (c-f) Dotplots representing expression level of cell cycle genes. The size of dots represents the percentage of cells with sequencing reads for each gene and color code represents average expression level. Expression level of G1/S and G2/M-specific genes are shown in (c) and (e), respectively, for 10X Genomics dataset with identical genes shown in (d) and (f) for Smart-Seq2 dataset. Individual dots are sized to reflect the proportion of cells of each type expressing the marker gene and colored to reflect the mean expression of each marker gene across all cells, as indicated in the key. (g and i) t-SNE plots labeled by estimated cell cycle status for 10X Genomics data (g) or Smart-Seq2 data (i). (h and j) stacked bar graph representing ratio of cell cycle stages in each cluster for 10X Genomics data (h) or Smart-Seq2 data (j). Embryonic cells were excluded from fresh cells and experimentally differentiated neurons were excluded from cultured cells in panel (j) because embryonic cells are highly proliferative unlike other fresh cells and differentiated neurons are post-mitotic cells unlike other cultured cells.

### Increased diversity of transcriptome consequential to *in vitro* expansion

Transcriptome alteration in cultured CPCs and meta-analysis of multiple cell types indicates a global (widespread) alteration of transcriptome rather than limited reprogramming of selected cell type-specific genes. Overall transcriptome diversity reflected by the number of detected genes in 10X Genomics dataset increased two-fold in cultured CPCs, corroborated by Smart-Seq2 dataset meta-analysis (median number of genes detected: 2,065 and 4,312 genes in fresh and cultured cells for 10X Genomics, and 3,490 and 6,528 genes in fresh or cultured cells for Smart-Seq2, respectively; Fig. 4a and 4b; p < 0.001, Wilcoxon rank sum test). Cultured cells still exhibit significantly increased detected genes after normalization by downsampling (median: 4,187 and 5,559 genes in fresh or cultured cells, respectively; Fig. 4c; p < 0.001, Wilcoxon rank sum test). These findings support the premise that *ex vivo* expansion of CPCs promotes increased transcriptome diversity despite loss of tissue-specific marker identity (Fig. 2 and 3).

**Figure 4.**
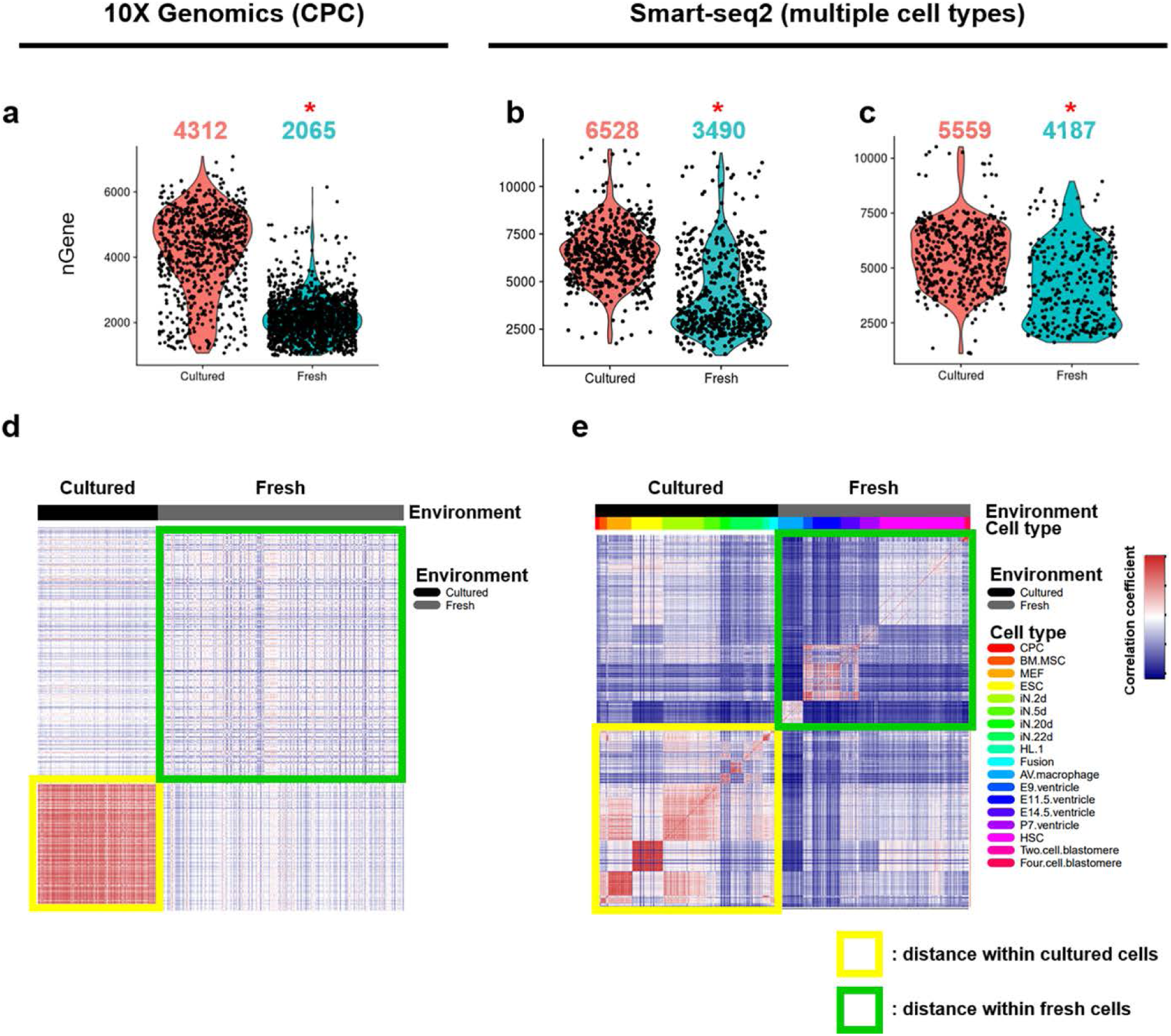
Cells acquire thousands of genes expressed during tissue culture while losing heterogeneity. (a – c) Violin plots represent the number of detected genes in freshly isolated cells or cultured cells in 10X Genomics (a) or in Smart-Seq2 (b and c) datasets. The number above the violin plots represent the median value of the number of detected genes. nGene, the number of detected genes; *, p-value < 2.2e-16, Wilcoxon rank sum test. (d – e) Heatmaps depicting similarity between single-cells, indicating cultured cells have higher similarity compared with the similarity between fresh cells in 10X Genomics (d) or in Smart-Seq2 datasets (e). CPC, cardiac progenitor cell; BM.MSC, bone marrow-derived mesenchymal stromal cell; MEF, mouse embryonic fibroblast; ESC, embryonic stem cell; iN, induced neuronal cell; HL-1, atrial cardiomyocyte cell line; Fusion, Fusion cell ofBM-MSC and HL-1; AV macrophage, atrioventricular node macrophage; E9, E11.5, E14.5, embryo 9, ll.5, 14.5 days, respectively; P7, postnatal 7 days; HSC, hematopoietic stern cell; nGene, the number of genes detected

Interestingly, transcriptome alterations prompted by *ex vivo* expansion increase population homogeneity compared to freshly isolated CPCs extracted from their native niche *in vivo* microenvironment (Fig. 4d). The postulate that cultured cells exhibit greater transcriptome similarity than freshly isolated cells was confirmed by Smart-Seq2 datasets (Fig. 4e). The environmental influence of cell culture promotes transcriptome migration toward a common shared profile throughout the population with minimal unique signatures. In summary, environment-dependent influences play a major role in determination of cellular transcriptome and *ex vivo* culture expansion of isolated CPCs dramatically alters their transcriptional profile.

## Discussion

Every cell type is characterized by a unique set of expressed genes known as identity markers together with commonly shared gene profiles. Retaining cellular identity is important to maintain functional properties that presumably are inextricably associated with biological activity. In the context of cardiac stem cell therapy, multiple cell types have been used for therapeutic intervention with standard protocols involving *in vitro* cell expansion to provide sufficient quantities for introduction into the damaged myocardium. However, drift in transcriptional identity of freshly isolated CPCs used for stem cell treatment consequential to *in vitro* expansion has not been addressed on the single cell level. Findings from this study derived from massively parallel digital transcriptomic profiling and meta-analysis of multiple scRNA-Seq datasets reveal that *in vitro* CPC expansion is associated with increased transcriptome diversity by acquiring expression of thousands of genes, up-regulation of cell cycle and metabolism genes, and loss of identity marker gene expression that increased transcriptional similarity among multiple types of cultured cells (Fig. 5). Transcriptional profiles in response to environment are largely conserved among various cell types, which has important implications for implementation and expectations of cardiac stem cell therapeutic implementation as discussed below.

**Figure 5.**
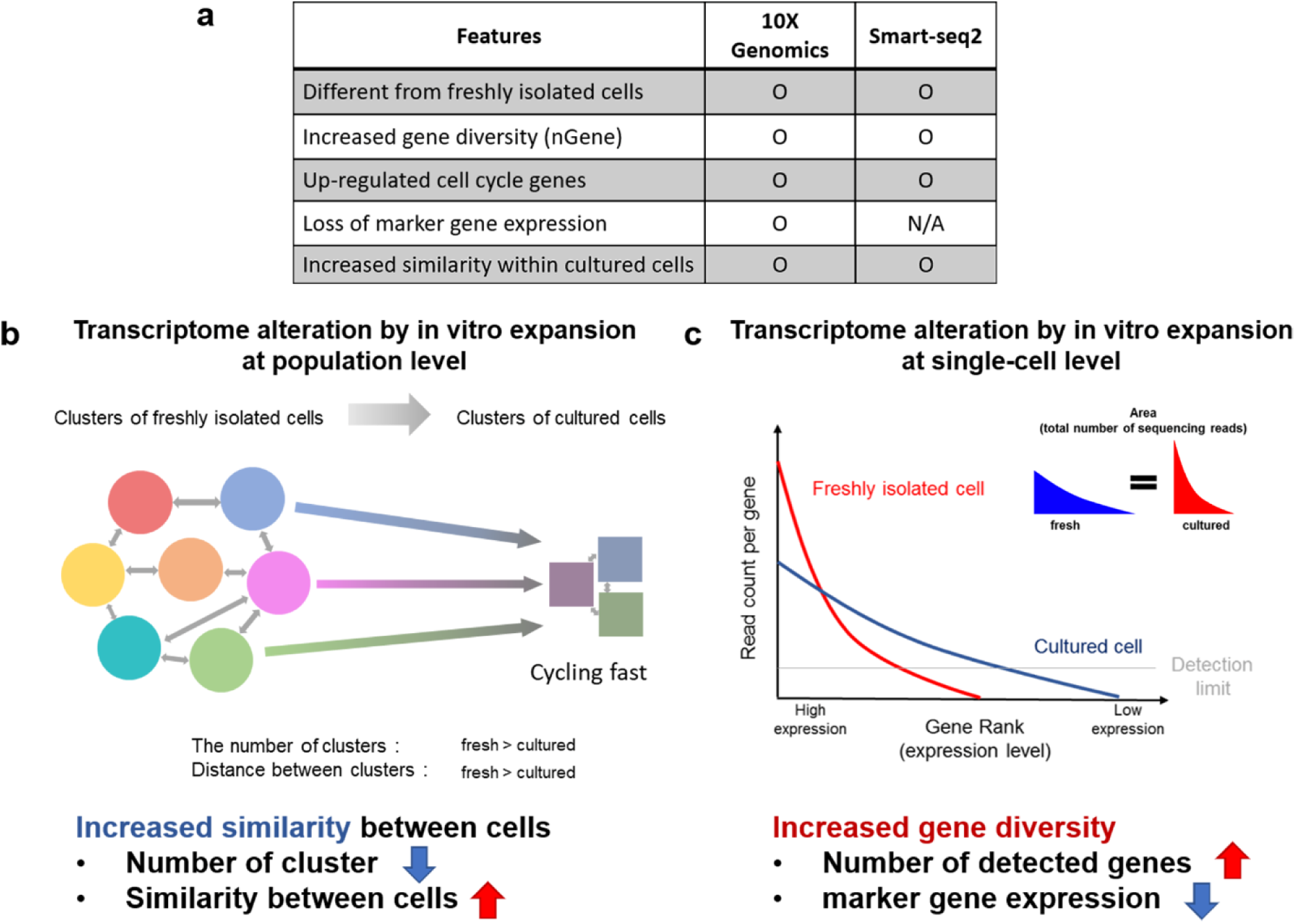
Transcriptome alteration by *in vitro* expansion. (a) Five key features of cells acquired during tissue culture. (b and c) Proposed model at the aspect of population (b) or single-cell (c) level. *Ex vivo* expansion in culture prompts loss of transcriptomic heterogeneity, leading to high similarity between cultured cells regardless of cell type. Expression of identity markers is suppressed with concomitant acquisition of expression for thousands of genes following culture characterized by increased transcriptome diversity.

Consequences and desirability of phenotypic and functional alterations to *ex vivo* expanded CPCs are largely unrecognized and uncharacterized, but there is ample evidence that culture conditions exert profound influence upon cellular biological properties.^31-34^ For example, low oxygen tension improved self-renewal and maintained differentiation potential^35^, whereas microcarrier and stirrer systems increased CPC expansion while retaining cellular phenotype and omics profiles.^31^ Stem cell immaturity and inhibition of gene methylation was promoted by replacement of serum with human supplements.^34,36,37^ Collectively, the intent of such studies is to improve cell expansion efficiency or the quality of the expanded cells through the manipulation of culture conditions, but the referential point of how such manipulations drift from ancestral fresh cell origins was not a factor in assessments. Consequences of *in vitro* expansion and biological drift upon cellular properties is an important consideration, especially if the intended outcome is to capitalize upon the identity and functionality of forerunners by creating an expanded derivative population with traits conserved from their origins.

Transcriptome profiles may be more profoundly altered by culture conditions depending upon cellular differentiation status. Culture conditions provoked relatively minor differences in mesenchymal stem cell (MSC) from uncultured predecessors at transcriptome level^12^, whereas terminally differentiated cells such as brain macrophages and endometrial cells are all altered substantially by *in vitro* conditions.^14,15^ Previous studies suggest CPC multipotentiality for lineage commitment^38^ that could lead to prediction of modest transcriptomic changes resulting from *in vitro* expansion as noted for MSC.^12^ However, results with CPCs presented in this report indicate profound changes in transcriptome with coordinated up-regulation of majority of cell cycle genes (Fig. 3), in contrast to down-regulated cell cycle inhibitors promoting increased cell cycle activity in MSC.^12^ Technical differences in transcriptome analyses could also play a role, as the microarray used for comparing fresh MSC with cultured MSC might not be sensitive as RNA-sequencing used for other cell types presented herein. Considerations of bulk versus scRNA-Seq and microarray versus whole transcriptome profiling will always present challenges when performing cross-platform comparisons of transcriptomic datasets and needs further attention to identify potential limitations and caveats.

The advent of scRNA-Seq technology has enabled a powerful advance in delineation of population heterogeneity of freshly isolated cells^17,39-41^. In comparison, relatively little attention has been focused upon characterization of population heterogeneity for donor cells prepared for cell therapy, particularly at the single cell level. The overarching findings presented herein show that CPC population heterogeneity decreases following *in vitro* expansion resulting in a more homogeneous transcriptomic profile. Future studies will need to determine the general applicability of these observations to additional cardiac-derived stem cell types as recently characterized by our group.^25^ Implications of donor cell population homogeneity for therapeutic stem cell treatment are significant, as the capacity of adoptively transferred effector cells to maintain their acquired *in vitro* transcriptomic profile or adapt to *in vivo* environmental conditions upon reintroduction to damaged myocardium remain wholly unexplored.

The substantial alteration of transcriptome as well as loss of original identity markers suggests a marked change in functional characteristics of *in vitro* expanded cells from their freshly isolated ancestors. A key question to be resolved remains as to whether alterations prompted by culture conditions exhibit plasticity upon introduction to intact tissue. Tissue culture influence upon efficacy of cell therapy and mitigation of undesirable transcriptional reprogramming requires systematic analyses using the multiple cell types currently being advances for clinical interventional approaches. Findings of considerable transcriptional drift and decreased population heterogeneity for *in vitro* expanded cells revealed in this study could well account for consistently modest outcomes of cardiovascular cell therapy regardless of chosen cell type.^42^ Greater appreciation of the impact, permanence, and functional benefits or impairments yielded by *in vitro* expansion of stem cells will contribute significantly toward development of improved protocols and cell preparations to enhance the reparative potential of adoptively transferred regeneration-associated cellular effectors.

## Methods

### Isolation of c-Kit^+^ CPC populations

Adult c-Kit+ CPCs were isolated and expanded as previously described^1^. Briefly, two FVB female mice hearts were perfused per group on a Langendorff system for blood removal, and tissue was subsequently digested for 10-15 minutes with Liberase DH digestion buffer (Roche 05401089001, 5 mg/mL in perfusion buffer) and dissociated through pipetting. After removing cardiomyocytes by cell strainers, Lin^-^ cKit^+^ CPCs were obtained by immunomagnetic sorting with Lineage depletion kit and CD117-conjugated Microbeads (Miltenyi Biotech 130-048-102). Fresh isolated CPCs were subjected to immediate single-cell RNA-Seq analysis or cultured for five passages and then used for single-cell RNA-seq. All experiments involving mice and use of vertebrate animals were carried out according to Institutional Review Boards (IRB) policy and approved by the Institutional Animal Care and Use Committee (IACUC) at San Diego State University.

### Single-cell RNA-seq

#### 10X Genomics platform

Freshly isolated and cultured CPCs suspensions were loaded on a Chromium^TM^ Controller (10x Genomics) and single-cell RNA-Seq libraries were prepared using Chromium^TM^ Single Cell 3’ Library & Gel Bead Kit v2 (10x Genomics) following manufacturer’s protocol. Each library was tested with Bioanalyzer (average library size: 450-490 bp). The sequencing libraries were quantified by quantitative PCR (KAPA Biosystems Library Quantification Kit for Illumina platforms P/N KK4824) and Qubit 3.0 with dsDNA HS Assay Kit (Thermo Fisher Scientific). Sequencing libraries were loaded at 2 pM on an Illumina HiSeq2500 with 2X75 paired-end kits using the following read length: 98 bp Read1, 8 bp i7 Index, and 26 bp Read2.

#### Smart-Seq2 platform

After trypsinizing cultured CPCs, single cells were captured under stereomicroscope by mouth pipetting with a ~0.2 mm diameter flame-pulled glass Pasteur pipet attached to aspirator tube (Sigma-Aldrich, A5177). Selected cells were dispensed into Eppendorf tube containing 10 μL cell lysis buffer provided by Smart-Seq v4 kit. cDNA was synthesized following manufacturer’s protocol (Smart-Seq v4 ultra low amount cDNA kit Clontech, 634888) and illumina sequencing libraries were then constructed using the Nextera XT DNA Sample Preparation kit (Illumina, FC-131-1024). The sequencing libraries were quantified by quantitative PCR (KAPA Biosystems Library Quantification Kit for Illumina platforms P/N KK4824) and Qubit 3.0 with dsDNA HS Assay Kit (Thermo Fisher Scientific). The pooled libraries were sequenced as paired-end 75×75 base reads on a NextSeq 500 with mid-output kit.

### Data Analysis

#### 10X Genomics platform

The raw data was processed with the Cell Ranger pipeline (10X Genomics; version 2.0). Sequencing reads were aligned to the mouse genome mm10. 1,615 and 850 cells were recovered from freshly isolated and cultured CPC samples, respectively. Cells with less than 1,000 genes or more than 10% of mitochondrial gene UMI count were filtered out and genes detected less than in three cells were filtered out.^2^ Altogether, 2,383 cells and 15,786 genes were kept for downstream analysis using Seurat R Package (v2.3.0). Approximately 2,000 variable genes were selected based on their expression and dispersion. The first 15 principal components were used for the t-SNE projection^3^ and unsupervised clustering^2^. Gene expression pathway analysis was performed using clusterProfiler^4^

#### Smart-Seq2 platform

Smart-Seq2 scRNA-Seq datasets were obtained from public databases (Supplementary table) with exception of the CPC dataset generated for this study. Sequencing reads were mapped to UCSC mouse genome mm10 using STAR v2.5.2b^5^ with default parameters and only uniquely mapped reads were kept. Read counts table was used as an input for generating Seurat object. Cells with less than 1,000 genes or more than 10% of mitochondrial gene count were filtered out and genes detected less than in three cells were filtered out. To exclude effects of sequencing depth variation, scRNA-Seq raw data were downsampled to 500,000 reads per cell by random selection of raw read from fastq files and re-analyzed. 18,698 genes and 1,126 cells were kept for further analysis. Clustering analysis and downstream analysis were performed as outlined in the 10X Genomics platform section.

## Statistics

Significant differences in the number of genes detected between fresh and cultured datasets from both platforms were analyzed with Wilcoxon matched-pairs rank sum test, meeting distribution assumption with statistical significance accepted when p < 0.05.

## Data availability

scRNA-Seq data generated in this study has been uploaded to the Gene Expression Omnibus (GEO) database (GSE114280). Seven public scRNA-Seq data sets were used in this study (see details in Supplementary table)

## Acknowledgements

The authors wish to thank all members of the Sussman lab for helpful input and valuable discussions.

## Author contributions

T.K. and M.A.S. designed the study. A.C. and K.M.B. isolated and cultured CPCs and B.J.W. captured single-CPCs for Smart-seq2. T.K. performed scRNA-Seq experiments and most of data analysis. B.-H.K. analyzed data. T.K. and M.A.S. wrote the article. O.H.E edited the article. All authors read and approved the final article.

